# Electrical signaling and coordinated behavior in the closest relative of animals

**DOI:** 10.1101/2024.06.19.599169

**Authors:** Jeffrey Colgren, Pawel Burkhardt

## Abstract

The transition between simple to complex multicellularity involves large degrees of division of labor and specialization of cell types. In animals, complex sensory motor systems are primarily built around the fundamental cell types of muscles and neurons, though the evolutionary origin of these cells, and their integration, remains unclear. Here, in order to investigate sensory-behavior coupling in the closest relatives of animals, we established a line of the choanoflagellate, *Salpingoeca rosetta*, which stably expresses the calcium indicator RGECO1. Using this, we identify a novel cellular behavior associated with electrical signaling, in which ciliary arrest is coupled with apical-basal contraction of the cell. This behavior, and the associated calcium transients, are synchronized in the multicellular state and result in coordinated ciliary arrest and colony wide contraction, suggesting information is spread amongst the cells. Our work reveals fundamental insights into how choanoflagellates sense and respond to their environment and offer a new perspective into the integration of cellular and organism wide behavior in the closest protistan relatives of animals.

## Introduction

Animals move through and interact with their environments in ways unique to life on earth. Calcium signaling is at the crux of this sensory-behavioral integration, playing a role in every modality from initial signal detection to downstream responses. This is achieved through dynamic control of free cytoplasmic calcium concentrations, and modulation of entry, release, and clearance. Though the emergence of calcium as a signaling molecule is very ancient, there are marked differences in the mechanisms employed by plants, fungi, and animals; the major lineages where complex multicellularity is present[1]. It is hypothesized that stable multicellularity provides strong pressure for the development of versatile communication systems between cells. In animals, sensory-motor integration and coordinated behavior is primarily achieved by the activity of specialized cell types, neurons and myocytes[2–4]. These are excitable cells which depend on large and tightly regulated calcium events to trigger a rapid response. On short timescales, this response can act to convert environmental information into electrical or chemical signals, propagate these signals, or convert the signal into force generation or movements. Based on their broad phylogenetic distribution, muscles and neurons likely arose very early in animal evolution and potentially multiple times through convergence[5–10]. Furthermore, many of the proteins involved in these pathways are also found outside of animals, with a large collection found in the closest unicellular relatives of animals[11–13]. However, the function of these components and their level of integration within these organisms largely remains unstudied.

The ability to precipitate phosphate ions makes high calcium concentrations inherently toxic to life as we know it, resulting in strong evolutionary pressure on sequestering, compartmentalizing, and extruding calcium ions[14–16]. In animal cells, cytoplasmic levels of free calcium are in the nM range, while extracellular concentrations are in the mM range, and major intercellular stores in uM range (up to mM range in excitable cells, such as muscles and neurons)[14–16]. Calcium is an important secondary messenger in animal excitable cells. As a general mechanism, membrane depolarization activates voltage gated calcium channels (VGCC) which leads to an influx in calcium. This influx can then trigger an immediate response (such a vesicle fusion in neurons) or further release of calcium stores (as in muscle contraction). These are transient events, and clearing of signal occurs via segregation by calcium binding proteins, compartmentalization into internal stores, and/or extrusion from the cell[17–19].

VGCC are broadly distributed in eukaryotes and have been shown to influence/regulate flagellar movements in paramecium and Chlamydomonas[20,21], mediate infection by various parasitic protists[22], as well as leaf movement in plants[23]. In animals, these channels can be broadly characterized into low-voltage activation and high-voltage activated channels[24]. Phylogenetic analyses has shown that choanoflagellates, the closest known living relative to animals, possess a single member of each of these categories[25].

Choanoflagellates are bacterivorous filter feeders and are highly polarized cells characterized by a single apical flagellum, surrounded by a microvilli-based collar (**Figure 1A**). The planar beating of the flagellum generates flow, which draws bacterial prey into contact with the collar where it is phagocytized **(Figure 1B)**. Choanoflagellates have complex life cycles, with some displaying transient multicellularity, which is generally achieved through serial cell division[26]. The choanoflagellate model *Salpingoeca rosetta* has at least 6 distinct cell stages; linear chain colonies, rosette colonies, slow swimming solitary cells, fast swimming solitary cells, attached cells and ameboid-like cells following physical confinement[27,28]. Both chain colonies and rosette colonies occur through serial cell division followed by incomplete abscission, which leaves a small cytoplasmic bridge between the cells[27,29] (**Figure 1C**). These bridges have only been observed by electron microscopy and contain two electron dense structures each localized near one of the cells. Under nutrient rich conditions, cells divide rapidly in chains before separating into slow swimmers. Environmental cues, such as the presence of specific bacterial sulfonolipids, can trigger the development of rosette colonies[30]. In this case, the cells secrete ECM from their base into the central region of the group of cells. The result is a multicellular ball with the collars and flagella facing outward. The cytoplasmic bridges are maintained in this state and can take on a highly complex arrangement[29].

**Figure 1.**
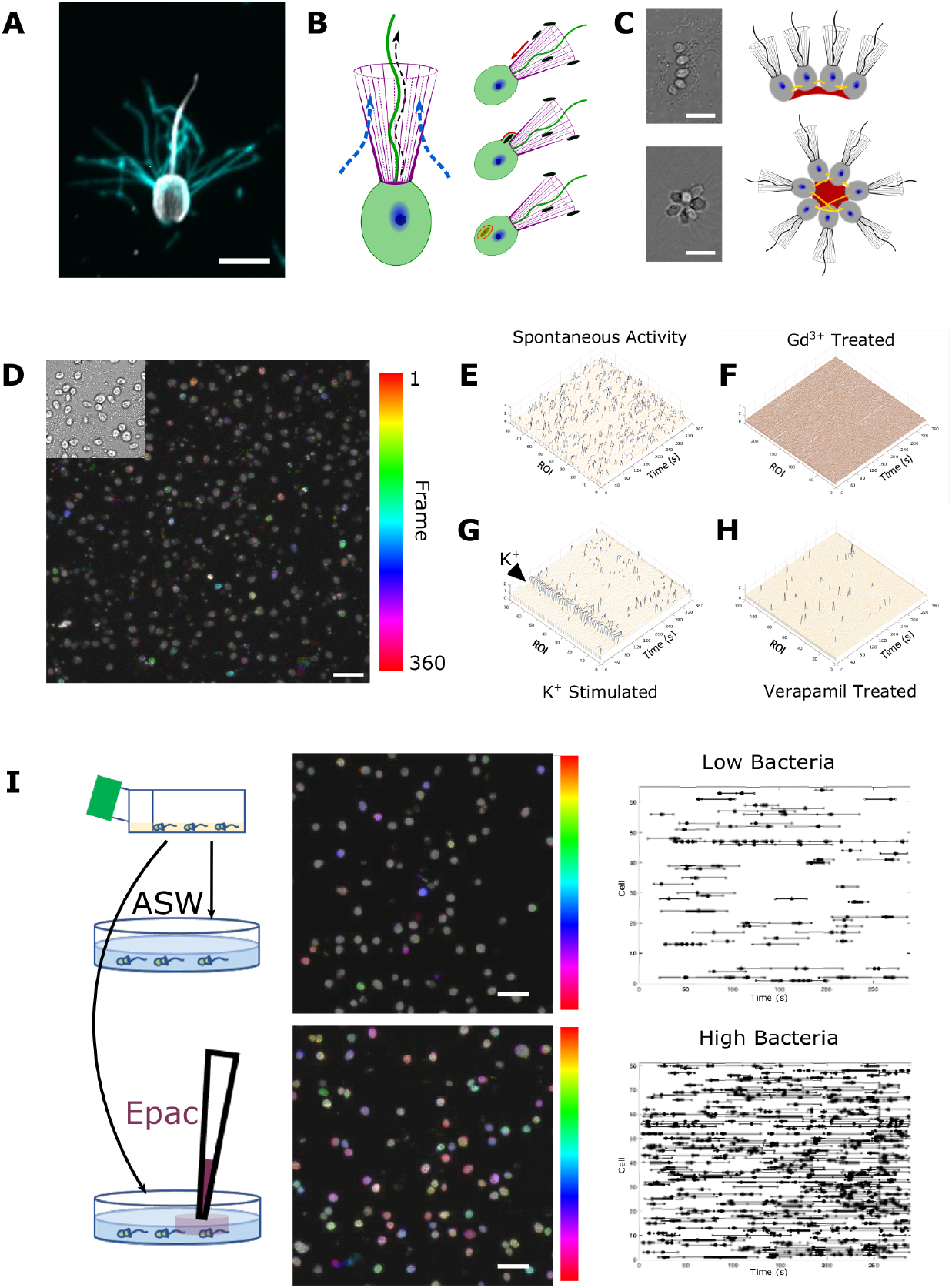
Morphology, feeding, and spontaneous Ca^2+^ dynamics. (A) Unicellular state of *Salipingoeca rose*.*a*, showing tubulin (grey) and actin filaments (magenta), the polarized cell has a single flagellum (cilia) surrounded by a microvilli based collar. **(B)** Planar beating of the flagellum draws water through the microvilli collar where bacterial prey can become trapped. Prey is phagocytized at the base of the collar **(C)** Multicellular states in *S. rose*.*a* form through serial division, resulting in chains (top) or rosettes (bottom). Cells are connected by thin intercellular bridges (yellow/orange in diagrams) and share ECM (red). In rosette filopodia (grey) at the base of the cells extend into a thick ECM orienting the collar and flagella outward. Intercellular bridges do not have a stereotyped pattern in colonies. **(D)** Time series projection of RGECO signal in cells over 360 frames (1fps) of a spontaneously acting culture. Upper leR shows a still of simultaneously acquired DIC images of the culture. **(E)** deltaF/Fo (z-axis) traces for all cells (y-axis) over time (x-axis) in a spontaneously acting culture. **(F)** A mirror culture treated with Gd^3+^ shows complete loss of spontaneous signal. **(G)** Perfusion of 25mM K^+^ (T=80s) triggers a highly synchronized event within the field of view. **(H)** Treatment with 10μm verapamil decreases spontaneous activity. **(I)** Cultures were thoroughly washed with artificial sea water (ASW) and then plated in ASW or ASW with a known quantity of feeder bacteria. LeR shows temporal projections of activity in a washed culture (top) and a mirror culture with bacterial prey reintroduced (bottom) prior to imaging. Right shows raster plots for each culture with dots indicating peaks and whiskers rise and fall (respectively). Scale bars 5μm (**A**) and 20μm (**C, D, I**).

Though morphologically simple compared to animals, choanoflagellates have been extremely successful, being found throughout the world’s oceans. There is evidence that the general morphology has existed for at least 600 million years, and it has been hypothesized that the first animal resembles a choanoflagellate colony[26,31,32]. Choanoflagellates have also been shown to demonstrate behaviors such as chemotaxis, aerotaxis, and pH-taxis[33,34], and have a large toolkit of ion channels and calcium binding proteins that function in animal sensory motor circuits[1,11,12,15,25].

Here we report on the calcium dynamics in the choanoflagellate *S. rosetta*, in both unicellular and multicellular stages. We find a range of spontaneous Ca^2+^transients, which can be grouped into discrete categories. We identify categories associated with VGCC activity and use this ‘calcium signaling vocabulary’ to identify a novel cell behavior. Furthermore, we find evidence of regulated flow of signal between the cells in colonies. We find that chains, the more transient form of multicellularity, show predominantly asynchronous events between cells, while the more stable rosette colonies demonstrate predominately synchronous events.

## Results

### Spontaneous calcium transients in single cells

To probe Ca^2+^ dynamics in *S. rosetta*, we established a line stably expressing the Ca^2+^ indicator RGECO (see methods and supplemental material for detailed workflow). Imaging of a culture in the unicellular stage revealed a large amount of spontaneous activity (**Figure 1D,E, & S1, Additional file 1**). This activity could be completely ablated by removal of extracellular calcium from the culture media (**Figure S2**). Perfusion of Ca^2+^ containing artificial sea water (ASW) showed a return of activity. The same results could be obtained by incubating cells in media containing the non-specific Ca^2+^ channel inhibitor gadolinium (5µM Gd^3+^) (**Figure 1F, Additional file 1**). Taken together, this suggests that the spontaneous activity is dependent on Ca^2+^ entry from the extracellular environment.

To understand activity of these cells, we perfused with depolarizing concentrations of K^+^ (25mM) while imaging and observed a highly synchronized response across the field of view (**Figure 1G, Additional file 1**). This could also be achieved with treatment of 10µM ATP (**Figure S3**). To validate that this response arises from cellular depolarization we fabricated a culture dish to allow for electric field stimulation (see methods) and applied sequential 10V pulses or a single 50V pulse (**Figure S3**). This resulted in a synchronized calcium transient across the culture followed by a return to asynchronous activity. Importantly, all cases of stimulation elicited transient events, while treatment with the calcium ionophore, Ionomyocin (1µM) showed a stable increase with limited bleaching of using the same acquisition settings (**Figure S4**). Since depolarization could elicit a consistent response, we treated with the VGCC inhibitor verapamil and found that it reduced, but did not completely ablate, spontaneous activity (**Figure 1H, Additional file 1**).

Next, to understand if there is any synchronicity within the spontaneous activity we performed network analysis on active cultures. The large number of transients showed a range of amplitudes as well as rise and fall times, with rising slightly faster than fall time (**Figure S5**). Analysis of pairwise synchronization identified 7 clusters of activity (**Figure S5**), while functional connectivity analysis did not identify major nodes of activity (**Figure S5**). Looking at the physical locations of individual cells within clusters suggests that the majority of synchronous activity is the result of random chance. However, clusters show some arrangement in space, suggesting activity may be tied to local environmental conditions (**Figure S5**).

Our choanoflagellate cultures are grown with *E. pacifica* feeder bacteria as the sole food source. We hypothesized that grouping of bacteria may be a determinant of the localized activity. To test this, cultures were extensively washed with artificial sea water to remove as much bacteria as possible. This culture was then split and plated without bacteria or with bacteria added back. In minimal bacteria cultures, spontaneous activity occurred at a rate of 0.027 events per min per cell, while cultures provided a large amount of bacteria showed activity at a rate of 0.356 events per min per cell (**Figure 1I & S6, Additional file 2**). This suggest that at least some of the activity is dependent on sensation of bacteria or feeding behavior.

### Voltage Gated Calcium Channel Dependent Transients

The *S. rosetta* genome contains two putative VGCCs (SrCa_v_1/2; PTSG_09464 and SrCa_v_3; PTSG_03773). To understand the contribution of VGCC in the spontaneous activity of cultures we treated them with increasing concentrations of the VGCC inhibitor verapamil and found a dose dependent decrease in spontaneous activity (**Figure 1H & S7, Additional file 1**). However, spontaneous activity was still present until lethal concentrations of verapamil were applied (**Figure S7**). This inability to mirror the complete loss of activity as with Gd^3+^ treatment suggests that some of the spontaneous activity relies on entry through verapamil insensitive channels. To extract information encoded in individual Ca^2+^ transients, we applied kShape clustering as a nonbiased means of grouping events. We clustered individual events within spontaneous acting cultures, K^+^ stimulated cultures, and verapamil treated cultures and identified 6 major classes of peaks (**Figure 2A**). By looking at how the proportions of these events changed in relation to different conditions we identified clusters 4 & 5 as likely being dependent on VGCCs, due to a decrease following verapamil treatment and an increase following stimulation (**Figure 2B**). Interestingly, these clusters also showed an increased proportion in the high bacteria culture (**Figure 2B)**.

**Figure 2.**
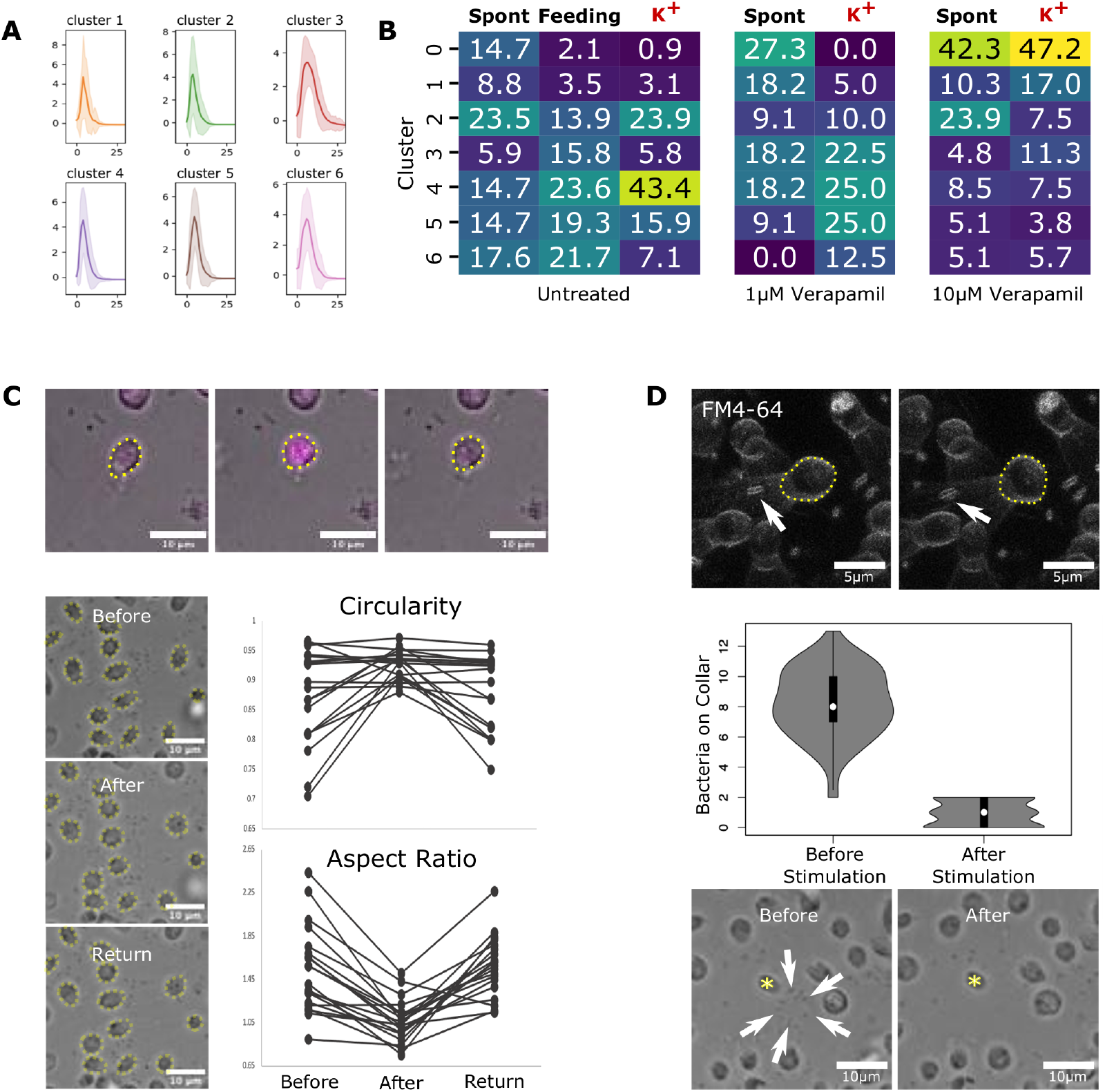
Events associated with VGCC activity are associated with a cellular behavior. **(A)** K-shape peak clusters from aligning individual peaks from spontaneously activity in normal culture, high bacterial, and verapamil treated, as well as from K^+^ stimulated. **(B)** Heatmap showing proportion of peaks that fall into each cluster in the different treatment conditions. Clusters 4 and 5 show the largest positive change to stimulation and negative change following verapamil treatment. **(C)** Top shows a DIC image overlayed with RGECO signal (magenta) of a cell undergoing a spontaneous event. The cell is outlined with yellow showing a shape change during the event. Bottom shows several cells in a field of undergoing a synchronized shape change following stimulation with K^+^. Quantification of circularity and aspect ratio of each cell shows a rounding up followed by return to original polarized shape during the event. **(D)** FM4-64 labeled cell (yellow outline) and bacterium (white arrow) at the collar, shows displacement of the bacteria during the shape change event. Below, quantification of visible bacteria on the cell’s collars before and just after stimulation. Still images show bacteria (white arrows) visible on the collar of a cell (yellow asterisk) before and after stimulation.

Next, we wanted to see if the different classes of transients could be correlated with any type of cellular behavior. Looking at DIC images of cells during class 4 & 5 events we found an obvious change in cell morphology, generally consisting of rounding up from apparent constriction along the apical-basal axis followed by a return to a more polarized shape during dissipation of the Ca^2+^ signal (**Figure 2C, Additional file 3**). This ‘behavior’ could be triggered by treatment with depolarizing concentrations of K^+^, concurrent with the synchronized transient event (**Figure 2C, Additional file 3**). These events can be generally characterized as a significant increase in circularity and decrease in aspect ratio (**P-values<0.001**) followed by a significant decrease in circularity and increase in aspect ratio (**P-values<0.001**). The duration of the constriction and relaxation closely mirrored the Ca^2+^ transients. During these events bacteria could also be seen being displaced from the feeding collar (**Figure 2D, Additional file 3**). Quantification of bacterial abundance before and after stimulation revealed a significant decrease in bacteria at the collar. Displacement of bacteria suggested that flow around the collar is altered during the event. Using high speed imaging, we watched flagellar movement during spontaneous and triggered events and found that the flagellum paused during the events, followed by a brief period of asynchronous beating, before returning to the normal pattern (**Figure 3A, Additional file 4**) and occurs in a synchronized manner following stimulation (**Figure 3B**). The initial rise in cytosolic Ca^2+^ levels preceded the flagellar arrest, suggesting the signal is propagated to the flagellum rather than from it (**Figure 3C & S8, Additional file 4**). High speed imaging of the cells undergoing flagellar arrest showed a straightening of the flagellum from the base towards the tip (**Figure 3D, Additional file 4**). For individual cells, the stereotyped sequence involved a cytoplasmic rise in Ca^2+^ followed closely (30-60ms) by flagellar arrest, which was followed by apical-basal contraction during the peak of [Ca^2+^]_cyto_. As RGECO signal returned to baseline, flagellar beating resumed prior to the return of cell shape (**Figure S9**)

**Figure 3.**
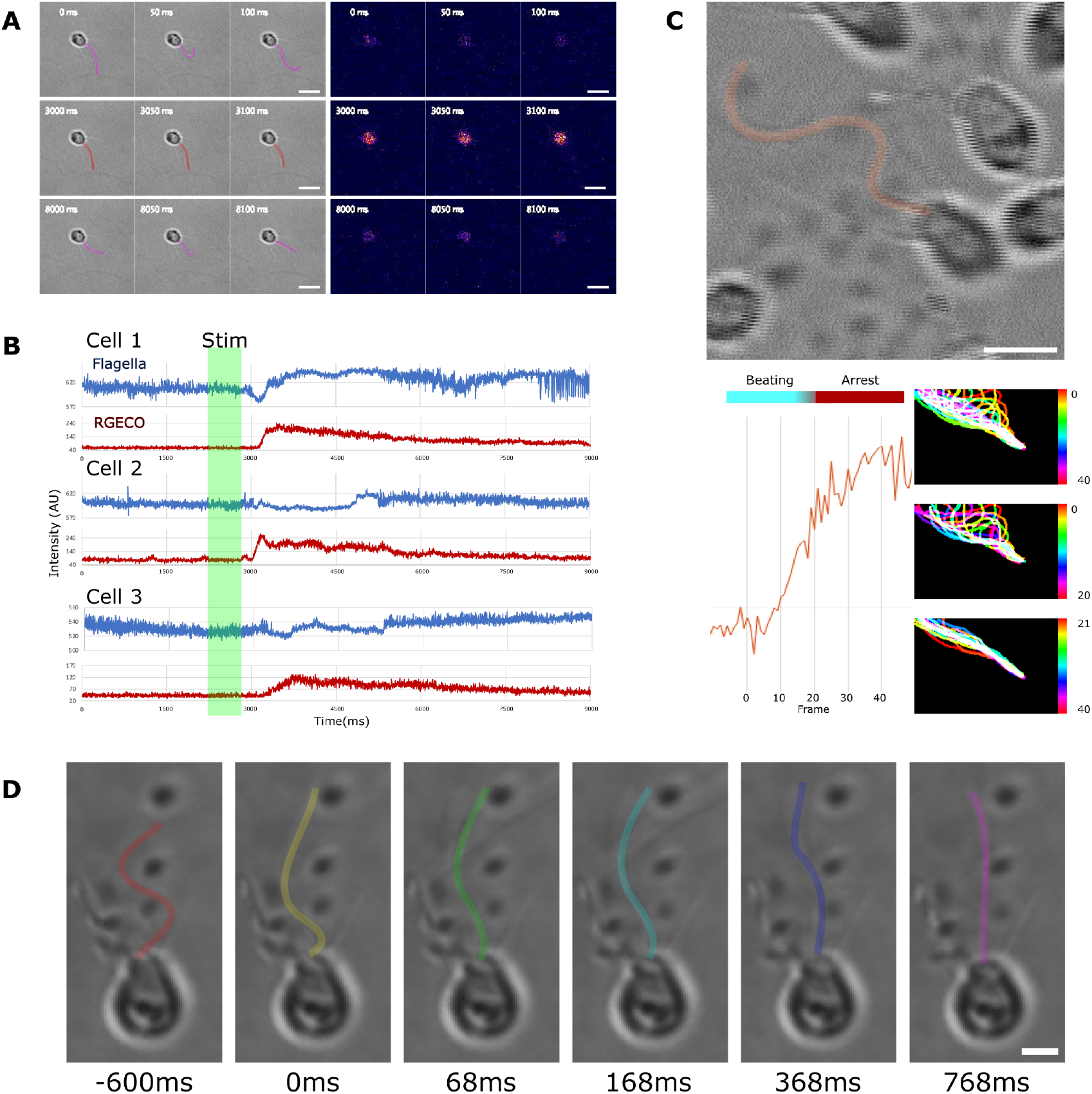
Calcium regulates flagellar arrest. **(A)** Flagellar beating arrests during events. Rows show 3 frames of overlapping DIC and RGECO images with the flagella highlighted in magenta (Top and bottom) or red (middle). Series occur before (top), during (middle), and after (bottom) and event. **(B)** Blue line shows pixel intensity of line scan taken parallel to the flagella originated from the base in the DIC images. Consistent beating shows periodic max and min values as the flagella passes the line, while changes in beating alter this pattern. Red line shows intensity of RGECO signal in the cell body. Green bar indicates when K^+^ was perfused into the culture. **(C)** Dual recording of RGECO signal and flagellar beating shows that arrest occurs after the initial increase in calcium concentration in the cell body. Bottom shows RGECO trace in ∼30ms frames on the left, with temporal projections of the flagella on the right. Top shows the projection over 40 frames during which RGECO signal reaches its maxima, the middle shows the first 20 frames, and the bottom shows the last 20 frames. **(D)** High speed imaging (568Hz) shows flagellar arrest occurs as a straightening from the base towards the tip. Scale bars 5μm (**A, C**) and 2μm (**D)**.

### Activity in Colonies

*S. rosetta* has at least 5 distinct cell stages including multicellular colonies consisting of either rosettes or linear chains[27]. These multicellular stages are inducible by environmental factors and form through serial cell division. To understand multicellular Ca^2+^ dynamics, we induced colony formation in our RGECO expressing line. In both rosettes and chain colonies we found that spontaneous activity occurred in a synchronized manner across the colony (**Figure 4A, Additional file 5**). As in single cells, spontaneous signals remained dependent on external Ca^2+^ availability and transients could be induced by depolarization. In chain colonies, the rapid depolarization of the full colony often led to separation of the cells. In spontaneously acting cultures, synchronized and asynchronous signals across the colonies were observed in both rosette and chain colonies (**Figure 4B**). Interestingly, the frequency of occurrence of each differed significantly between the two stages. In rosette colonies, synchronized signals were most frequent, making up **62.93%** of the events. Non-maximal synchronized events (characterized as at least one cell, but not every cell) occurred at **33.6%**. However, in **3.45%** of active rosettes, colonies showed rapid (within 3 seconds) asynchronous events between the component cells (**Figure 4A, Additional file 5**). These showed no pattern within spread around the colony. (**Figure 4A, Additional file 5**). On the other hand, asynchronous events seemed to dominate the spontaneous activity in chains, comprising **77.88%** of observed events. Synchronization within a subset of the colony was observed **8.85%** of the time, while synchronous signal across the colony was observed **13.27%** of the time (**Figure 4A&B, Additional file 5**).

**Figure 4.**
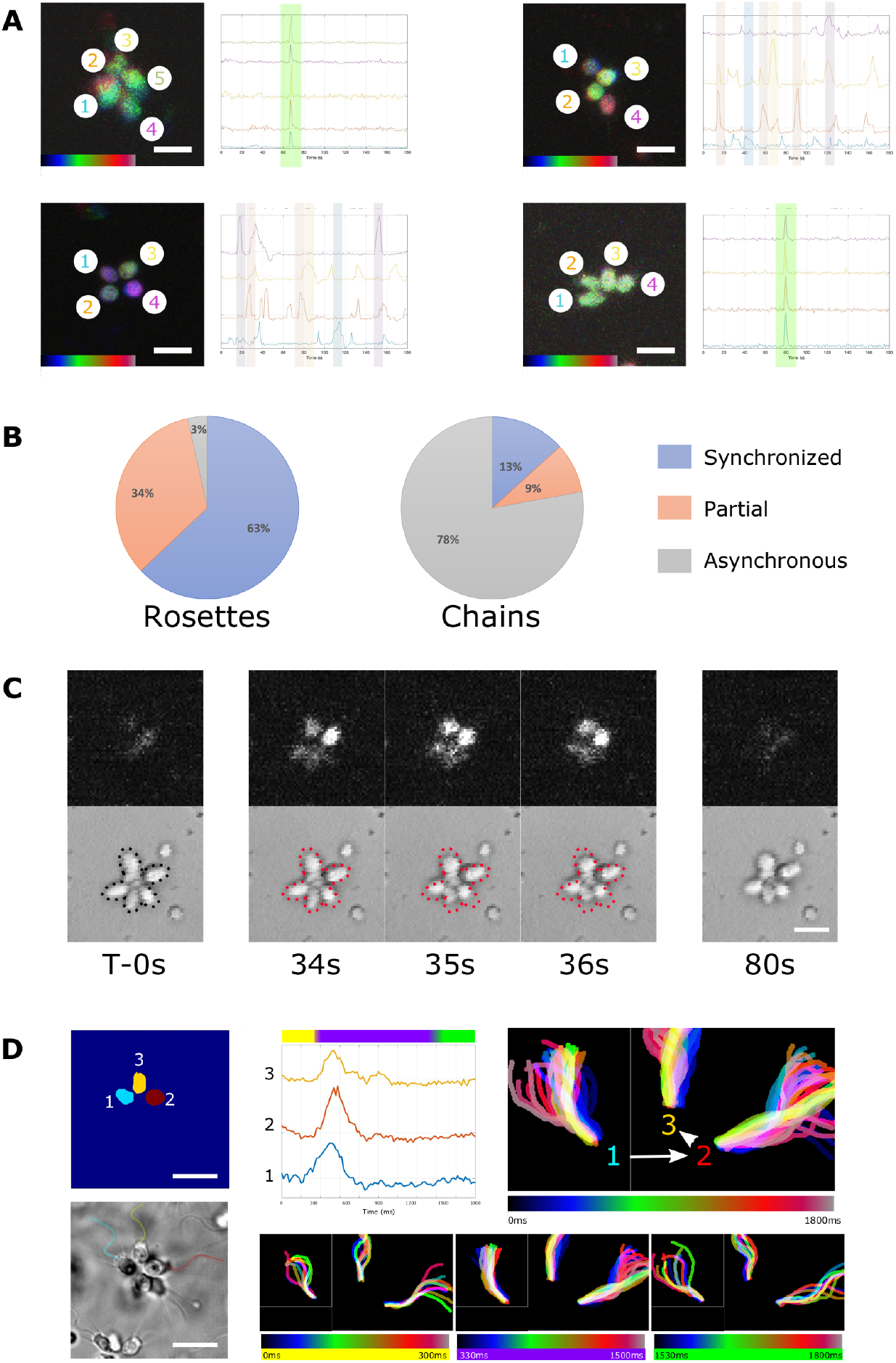
Both synchronized and asynchronized events occur in colonies and newly described behavior is synchronized. **(A)** Rosette colonies primarily show highly synchronized (top) but do have asynchronous events (bottom). Chains primarily show asynchronous events (top) while synchronous ones are present. Graphs show RGECO intensities for each cell, with color matching the number in the temporal projections. **(B)** Quantification of the occurrence of synchronized, partially synchronized (two or more cells) and asynchronous (multiple cells at different times) in spontaneously acting cultures. **(C)** REGCO (top) and DIC (bottom) images before, during, and after an event, show coordination of signal and contraction in the colony. **(D)** High speed imaging of spontaneous activity in a rosette shows synchronized flagellar arrest as well. RGECO signal first increases in cell 1, before visibly increasing in cells 2 and 3. All three visible flagella stop beating around 60-90ms after signal increase, and show a synchronized arrest. Temporal projections of flagella show the total time series, with the bottom panels showing the first 300ms during which cytoplasmic calcium increases, the ∼1200ms that beating is arrested, and the recovery period. Scale bars 10μm.

Strong transient events associated with the behavior described in single cells were observed in colonies and depolarization of the cells triggered a similar response. In rosette colonies, this led to a contraction of the full colony (**Figure 4C, Additional file 6**). Also like the single cells, the flagellar beating would briefly arrest (**Figure 4D & S10, Additional file 6**) during the event and bacterial shedding could be observed (**Figure S11**). When these events occurred, they were synchronized across the colony. Cytoplasmic Ca^2+^ levels began to rise prior to flagellar arrest, with similar kinetics, and spread of signal could be observed between the cells within the focal plane for which flagella were also visible (**Figure 4D, Additional file 6**).

## Discussion

One of the fundamental questions of the origins of complex multicellularity is how isolated cells coalesce into larger assemblages capable of specialization and coordinated behavior. Our observation of both synchronized and asynchronous transients in colonies suggest that regulated signaling occurs between these cells. Cells in colonies are linked by thin cytoplasmic bridges, believed to result from incomplete abscission[27,29]. The molecular composition of the cytoplasmic bridges between cells remains elusive. However, the tight synchronization observed in rosettes suggests that these connections are at least permeable to ions. Interestingly, the observed asynchronous activity within colonies further suggests that there is not open flow of ions between cells, rather a regulated process. The striking disparity in frequency of synchronized events in rosette colonies and chain colonies suggests that there is a fundamental difference in these two multicellular states. Increased synchronization is consistent with communication and coordination across colony, which allows it to act more as an individual unit as opposed to a collective. It is hypothesized that rosette colony formation primarily acts as a means of increasing feeding efficiency in food rich environments or as a means of increasing size to decrease predation[35–40]. While these are not mutually exclusive, and have both been documented in different instances of simple multicellularity, evidence suggests it does not lower rates of predation in choanoflagellates[41]. Though the contraction and flagellar arrest behavior occurs in colonies, it does so at a reduced rate, which is consistent with a higher holding capacity in the multicellular stage compared to the unicellular stage. Though further studies will need to be done, our findings are consistent with rosette formation functioning optimizing feeding efficiency.

In extant glass sponges, animals lacking both muscles and neurons, feeding is performed by drawing water through their body, via the beating of flagella on choanocytes, while food particles are trapped and phagocytized at the microvilli collar. These animals are composed of syncytial tissue which propagates action potentials, resulting in the coordinated and transient arrest of flagellar beating[42]. This coordinated behavior is thought to regulate the flow rate through their aquiferous system in order to maximize feeding efficiency[42–44]. Though the striking similarity to what we have found in choanoflagellate colonies could have arisen by convergent means, if this dates back to the last common ancestor of choanoflagellates and sponges, then it likely played an important role in animal multicellularity. Our findings suggest an important role of electrical signaling feeding behavior for *S. rosetta*. Though we find instances of phagocytic cups forming during strong calcium transients (**Figure S12**), the primary response appears to consist of the arrest of the flagella, contraction along the apical-basal axis, and shedding of bacteria from the feeding collar. Modeling of choanoflagellate filter feeding suggests that the bacterial load on the collar strongly affects the ability of the flagella to generate flow as well as the cells’ ability to propel itself forward[45,46]. It is possible that the here described behavior functions to balance feeding efficiency with the ability to disperse or simply maintain a high level of flow through the collar region. Choanoflagellates primarily interact with their environment by drawing it towards the cell through flagellar beating[25,47]. Though raised in the lab with a signal bacterial food source, the environments and microenvironments that these cells live in are not static and can contain bacterial blooms and swings of nutrient availability, as well as predators. This creates strong selective pressure for tight regulation of this process.

The capacity to move through and rapidly respond to a changing environment is a hallmark of animal biology. It has been hypothesized that early animal motility relied on coordinated regulation of ciliary beating across the organism[48,49] and likely was regulated by Ca^2+^[11]. However, much of this likely occurred during the time between the first obligately multicellular animal and the last common ancestor of extant lineages. The ability to directly sense and respond to gradients of stimuli in the environment is fundamentally constrained by a cells size[50,51]. As microeukaryotes, *S. rosetta* is approaching the lower limit on spatial sensitivity in its unicellular state. In rosettes, the size of the colony is sufficient to spatially resolve gradients, though prior studies have shown no coordination in the flagellar beating[52]. Here, for the first time, we find evidence that information flows between cells of a choanoflagellate colony in a regulated manner. Action potential-like control of ciliary activity has been reported in a variety of protists, often involved in facilitating and modifying movement[48,49,53]. We find the stereotyped response of ciliary arrest coupled with apical-basal contraction of the cell is synchronized with signaling across rosettes, culminating in a coordinated behavior. Information spread and coordination across a colony/mass of cells allows this to act as a single unit, meaning cells sample the environments across the full surface area, without the need to directly contact it. This is an essential step to open evolutionary space for modularization and sub-functionalization of cell states and may have helped provide the platform for complex multicellularity to arise in animals.

## Materials and Methods

### *Salpingoeca rosetta* cell culture and colony induction

*S. rosetta* (ATCC: PRA-390) was cultured with a single bacterium *Echinicola pacifica* as described in Booth, *et al*. 2018. Freezer stocks were thawed and seeded into 0.2x high nutrient media (HNM)[54] and grown for 3-5 days. Cultures were then maintained in HNM at 22°C and 60% relative humidify and were passaged at 1:60 every two days. Rosettes were induced by inoculating cultures with *Algoriphagus machipongnensis*.

### Plasmid design

RGECO1 was codon optimized for *S. rosetta* using highly expressed transcripts[55]. Codon optimization was performed with OPTIMIZER[55]. gBlocks fragments (Integrated DNA Technologies) were ordered with overhangs to facilitate HiFI assembly (New England Biolabs; E2621). The codon optimized RGECO1 was cloned into the *S. rosetta* expression plasmid NK802 (Addgene; #166056)[28] replacing the mCherry sequence. The assembled plasmid was transformed into NEB 5-alpha (high efficiency) competent cells (New England Biolabs; C2987). After verifying the sequence, the plasmid was purified using QIAGEN Endofree plasmid Maxi kit (QIAGEN; 12362). Purified plasmid was concentrated to 5mg/mL.

### Transfection

Transfection of *S. rosetta* was performed as described in Booth, *et al*., 2018. Prior to transfection cells were seeded at 8000 cells/mL in 40mL of HNM in a 175mL culture flask (Sardstedt; 83.3912.502) and grown for 48hr. On the day of transfection, cells were washed 3x with artificial sea water (400 mM NaCl, 8 mM KCl, 14.8 mM MgCl_2_*6H_2_O, 20.3 mM MgSO_4_*7H_2_O, 2.7 mM CaCl_2_*2H_2_O, 2.38 mM NaHCO_3_, 10 µM NaBr, 97 µM H_3_BO_3_, 100 µM SrCl_2_*6H_2_O, 10 µM NaF, 0.2 µM Kl) by pelleting at 2000xg for 5 minutes at room temperature (2200xg for the last spin). Cells were then resuspended at a concentration of 5×10_7_ cells/mL and 5×10_6_ cells were pelleted at 800xg for 5 minutes at room temperature. Supernatant was removed and cells were resuspended in 100μL of priming buffer (40 mM HEPES-KOH, pH 7.5, 34 mM lithium citrate, 50 mM cysteine, 15% PEG 8000, 3 μM papain), and incubated at room temperature for 35 minutes to digest extracellular matrix. Priming was halted by addition of 10μL of 10% BSA and the cells were pelleted at 1200xg for 5 minutes at room temperature. Primed cells were resuspended in 25μL of SF-Buffer (Lonza) and placed on ice. The transfection mixture was made of 14μL ice cold SF buffer, 2μL of SrRGECO plasmid, and 2 μL of carrier plasmid (pUC19 at 20ug/uL). Just prior to transfection 2μL of primed cells were added and the full volume was transferred to a well of a nucleofection cuvette (SF Cell Line 4D X Kit S (32 RCT), Lonza). A CM-156 pulse was applied in the 4D-Nucleofector (Lonza). 100μL of ice cold recovery buffer (10 mM HEPES-KOH, pH 7.5, 0.9 M sorbitol, 8% Polyethylene glycol 8000) was immediately added to the well and gently mixed by tapping on the side. After 5 minutes the full contents of the well were transferred to 2mL of low nutrient media in a 6 well plate and incubated at 22oC and 60% relative humidity for 30 minutes for cells to recover. 10uL of 10mg/mL *E. pacifica* was then added to the well and gently mixed. Transfected cells were then incubated at 22oC and 60% humidity.

### Selection for Stable integration

Following transfection, cells were incubated for 24 hours at 22oC and 60% humidity. After 24 hours selection began by addition of puromycin to a final concentration of 40μg/mL and cells were incubated 72 hours at 22oC and 60% humidity and monitored for growth. The upper 1mL of cells, which contained swimming chain colonies, was removed and pelleted at 2000xg for 10 minutes at room temperature. The cells were then resuspended in 2mL of HNM containing 80μg/mL puromycin and incubated at 22oC and 60% humidity for 48 hours. After incubation, washing was performed again but cells were resuspended in 12mL of HNM+80μg/mL and plated across a 6-well plate (Thermo Scientific; 140685). Cells were then incubated for 72 hours at 22oC and 60% humidity. Wells were monitored for growth and washing steps were repeated 2 times for actively growing cultures. Once consistent growth was observed cultures were passed 1:10 into fresh selection media and incubated for 48 hours at 22°C and 60% humidity. This was repeated two times. Small aliquots of each culture were then taken and the presence of fluorescence was confirmed by addition of 100μg/mL ionomycin (Milipore; 407950). Cultures which showed RGECO1 signal were then diluted to 3 cell/mL in selection media and plated in 100μL volumes on 96 well plates. Plates were then incubated for 96 hours at 22oC and 60% humidity and monitored for growth. Wells showing growth were marked and passed 1:10 into fresh selection media in 96-well plates. This was repeated two times after which 58 actively growing monoclonal cultures had been isolated. These cultures were screened for fluorescence signal by mixing 1μL of culture with 1μL of Ionomycin solution (100μg/mL final concentration) and viewing through an RFP filter. 26 of these cultures, which showed strong even signal, were then passed (1:10) into 300μL of selection media in a 48-well plate and incubated for 48 hours at 22oC and 60% humidity. After growing to density, 100μL of each of these cultures was screened live and during perfusion of ionomycin (100μg/mL final concentration) in order to judge the dynamic range of the sensor. Cultures which showed low baseline signal but strong maximal signal were then scaled up and frozen in 1:10 DMSO overnight in a Mr. Frosty freezing container (ThermoFisher – 5100-0001) at -80oC and then stored in liquid nitrogen. To establish running cultures, vials were thawed at room temperature and then added (1:10) to low nutrient media containing 40μg/mL puromycin. Growth was monitored alongside wildtype cultures which were not under selection (**Figure S13**)

### Plating and imaging

General imaging was performed on running cultures at a density of ∼106 cells/mL that were between passage 4-15. Glass bottom dishes (MatTek – P35G-1.5-10-C) were prepared by adding a drop of poly-L-lysine solution (0.01%) (Sigma-Aldrich - A-005-C) and incubating at room temperature for 30 minutes. Following incubation, the glass was washed 3x with ASW. Meanwhile, 1mL of cell culture was washed of excess bacteria by pelleting at 1500xg for 10 minutes at 4oC and the supernatant was carefully removed (using a gel loading tip for the final amount). Cells were then washed and re-pelleted in 1mL of ASW. The washed pellet was then resuspended in 100μL of ASW and transferred to the microwell of the glass bottom dish and incubated for 30 minutes at room temperature. Following incubation an additional 100μL of ASW was gently pipetted into the microwell to overfill it and the rest of the dish was filled with 2mL ASW without disturbing the well and cells were imaged.

### Imaging

Imaging was primarily performed on an Olympus FV3000RS confocal microscope (Olympus) using FLUOVIEW acquisition software (Olympus). General culture activity was aquired using a 60x objective (Olympus; UPlanSApo 60x/1.30 Sil) which contained ∼200 cells per field of view under normal plating conditions. Videos were acquired using a resonance scanner. RGECO1 was detected using RFP filter settings (530-550 excitation/575-625 emission) with transmitted light channel aquired through a DIC prism. 546 LED laser level was set between 2-3.5% and pinhole aperture was opened to balance speed and voltage was adjusted under hi-lo lookup table to a level below saturation in all visible cells. Stacks were set to a range just below and just above the cells and optimized based on aperture size. The same acquisition settings were used for all samples in an imaging session as well as sampled compared between sessions. Flagellar beating videos were acquired as a single optical section using a 100x objective (Olympus; UPlanSApo 100x/1.40 Oil) with the pinhole fully opened using the resonance scanning setting. Perfusions were performed by manually pipetting (at times supported by a stabilizing stand and micromanipulator) a concentrated solution into the solution outside the microwell to minimize changes in flow over the cells and allow time for diffusion to occur prior to interacting with them.

### Image analysis

When stacks were acquired, summed projections were generated in FIJI[56] and RGECO and DIC channels were separated. Primary analysis was performed using either FluoroSNAPP[57] or Mesmerize[58] analysis platforms (**Figure S14**) as well as PeakCaller[59] for visualization of cell traces and basic features. In FluoroSNNAP image series were checked for motion artifacts and aligned if needed. Segmentation was performed using active contour over time averaged series. Segmentation files were manually checked against DIC images and ROIs were removed if needed. For high frame rate series (ciliary beating) ROIs were manually drawn for cells chosen based on orientation of beating plane. Baseline fluorescence was set of a 10 second window and 50 percentile. Functional connectivity was calculated with cross-correlation (N resampling 100, Max time lag 0.5s), partial correlation (alpha-level significance 0.001), instantaneous phase (N resampling 100, alpha-level significance 0.001), Granger causality (max model order 20, alpha-level 0.05, with # iteration automatically detected), and transfer entropy (N resampling 100, Lags 1:10). For Mesmerize[60] analysis, motion correction and was performed using CaImAN[61] NoRMCorre[62] and ROIs were detected with CNMF(E)[63]. Samples were then normalized and deltaF/F0 was extracted and z-scored. Smoothing was employed using a Butterworth filter. Automated peak detection was performed and then manually checked for artifacts. Peak features were then extracted. k-Shape clustering[64] was performed on deltaF/F0 peak-curves with the data divided into partitions between 4 and 8, created by sorting the peak curves according to peak-width, with 100 different centroid seeds for each partition. 100 iterations of the kShape clustering algorithm were performed with a tolerance of 1e-6 and training performed on 30% of the data (**Figure S15**). For PeakCaller[65] visualization, extracted fluorescence from established ROIs was loaded. Parameters used were required rise, 60%; max lookback pts, 25; required fall, 60%; max lookahead pts, 25; over absolute %; trend control, finite difference diffusion (2-sided); trend smoothness 60.

DIC images were analyzed in FIJI[56]. For ciliary beating analysis, stacks were opened in LabKit plug-in[66] and manually annotated. Annotations were then opened in ImageJ and converted to binary masks. Masked files were converted to hyperstacks and temporal projections were generated over the full stack or on substacks within it.

### Functional Imaging

Calcium free media (CFM) was prepared by excluding Ca^2+^ from artificial sea water and including 10mM EDTA. During plating, cells were washed into and maintained in CFM. In order to transfer cells to normal media, the upper volume of CFM was removed from the dish and replaced with high Ca^2+^ media (10mM) and this was repeated with normal ASW. For gadolinium treatment, plated cells were mixed 1:1 with ASW containing 10μM Gd^3+^ and then the dish was filled with ASW+5μM Gd^3+^. For verapamil treatments, 2x verapamil solution in ASW was added to plated cells after 20 minutes and they were incubated for an additional 10 minutes prior to imaging. For ionomycin treatments, stock solution (10mg/mL in DMSO) was diluted to 2x concentration in ASW and mixed 1:1 with cells. For perfusion assays, a 10x ionomycin stock was gently pipetted in for a final 1x concentration. For electric field stimulation, a stimulation chamber was fabricated in a glass bottom dish. Wire channels were cut using a rotary saw and perpendicular platinum wires were fed through ∼1mm apart and affixed with silicone tape. A coverslip fixed over the middle and cells were plated beneath this, between the electrodes. All wires were stabilized by rigid supports on the microscope table and pulses were delivered manually with an isolated stimulator (npi – ISO-STIM_01D). For bacterial abundance assays, 40 mL of culture was collected and pelleted at 1500xg for 5 minutes at room temperature, and the supernatant was discarded. The cells were then resuspended in 50mL of ASW and shaken thoroughly before pelleting again. This was repeated 3x. The washed pellet was then resuspended in 400μL of ASW and an aliquot was taken to check cell density. The cells were then diluted to better match previous experiments and plated. To replace washed bacteria, a 10mg pellet of *E. pacifica* bacteria was resuspended in 500μL of ASW. For ‘low bacteria’ the cells were supplemented with 100μL of ASW and for ‘high bacteria’ cells were supplemented with 100μL of containing 2μg of *E. pacifica*. Intermediate bacterial concentrations were obtained by diluting 20μg/mL stock in ASW.

## Data and Materials Availability

All data has been deposited in FigShare (DOI:10.6084/m9.figshare.26065234). SrRGECO1 expression plasmid is deposited at Addgene (ID 222940) and SrRGECO cell line is available upon reasonable request.

## Author Contributions

J.C. and P.B. conceptualized and designed the research. J.C. generated the choanoflagellate line.

J.C. collected and analyzed imaging data. J.C. and P.B. wrote the manuscript.

## Acknowledgements

We thank all members of the Burkhardt group, past and present, for their thoughtful discussions. We would also like to particularly thank Aishwarya Ravi and Ronja Göhde for their assistance with the choanoflagellate cultures. We thank Marios Chatzigeorgiou and Timothy Lynagh for their conceptual and technical assistance with this work and we thank Scott Nichols for his valuable feedback on the manuscript.

