## Supplementary Information for "Electrical signaling and coordinated behavior in the closest relative of animals"

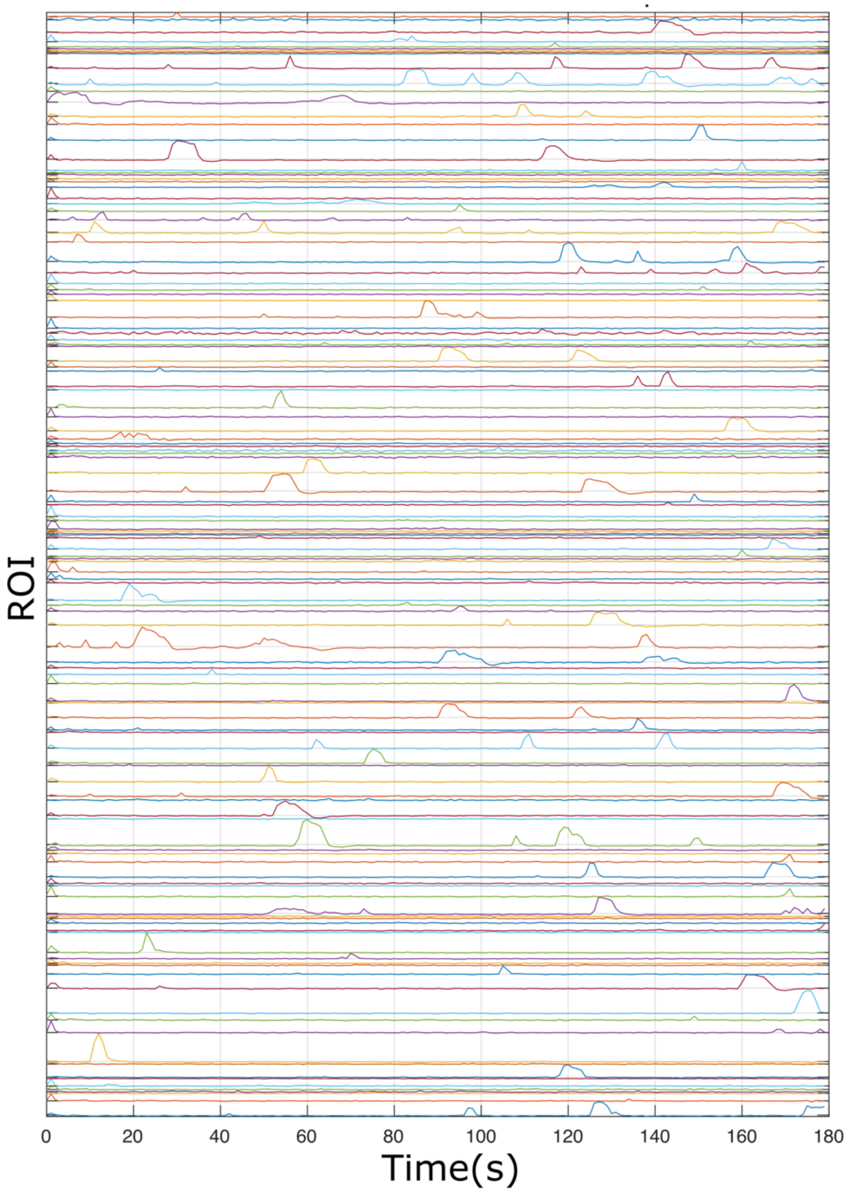


**Figure S1. Individual traces from spontaneously acting culture.** ΔF/F over time for individual cells in a field of view (ROIs) in a spontaneously acting culture. Cells were prepared by removing media and excess bacteria and washing once in clean ASW prior to plating.

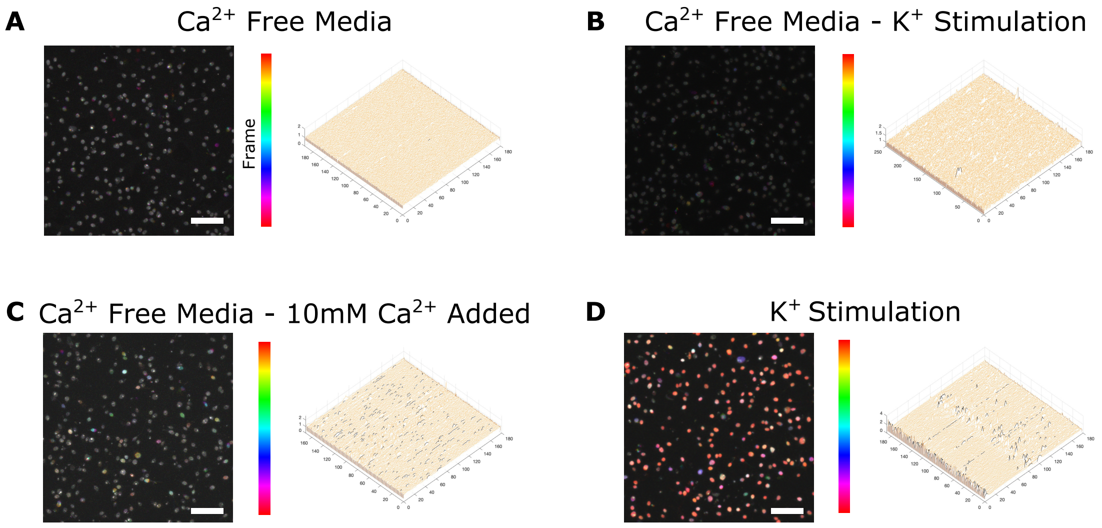


**Figure S2. Removal of External $Ca^{2+}$ ablates spontaneous activity. (A)** Incubation of cells in CFM shows a complete loss of spontaneous transients. **(B)** Stimulation with depolarizing concentrations of $K^{+}$ does not elicit a response in the absence of external $Ca^{2+}$. **(C)** Perfusion of $Ca^{2+}$ back into the media resulted in a return of spontaneous activity. **(D)** Cultures could then be stimulated by depolarizing concentrations of $K^{+}$. Temporal projections generated from videos in Supplemental video 1**.** Scale bars 50µm.

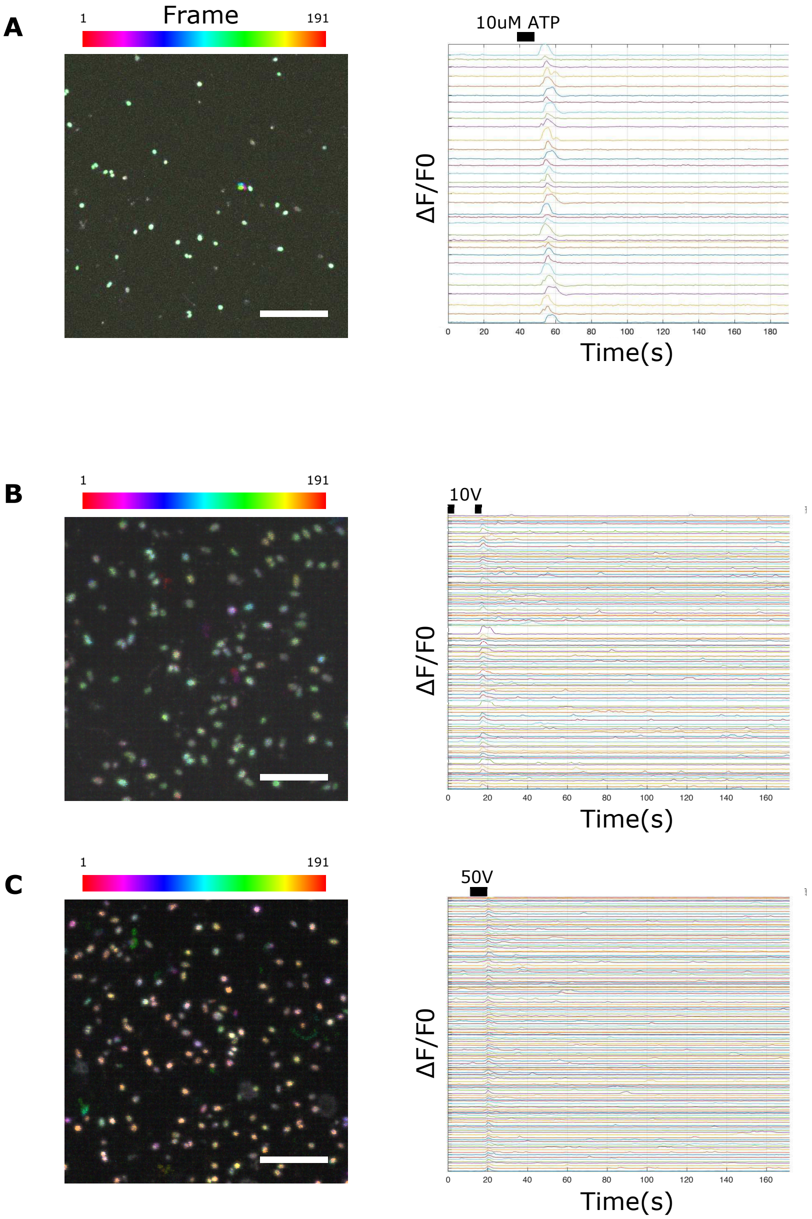


**Figure S3. Stimulation synchronized response. (A)** Left shows a temporal projection of a culture during perfusion with media containing 10uM ATP. Right shows individual traces for ROIs in the field of view, showing a strong and synchronized peak following application of ATP. **(B)** Cells were plated between electrodes in an electric field stimulator and manually pulsed with either 10V (top) or 50V (bottom). Temporal projections (left) show large amount of spontaneous activity still occurs in both cultures, while individual traces (right) show a synchronized peak following stimulation. Temporal projections and traces are generated from Supplemental video 2. Scale bars 50µm.

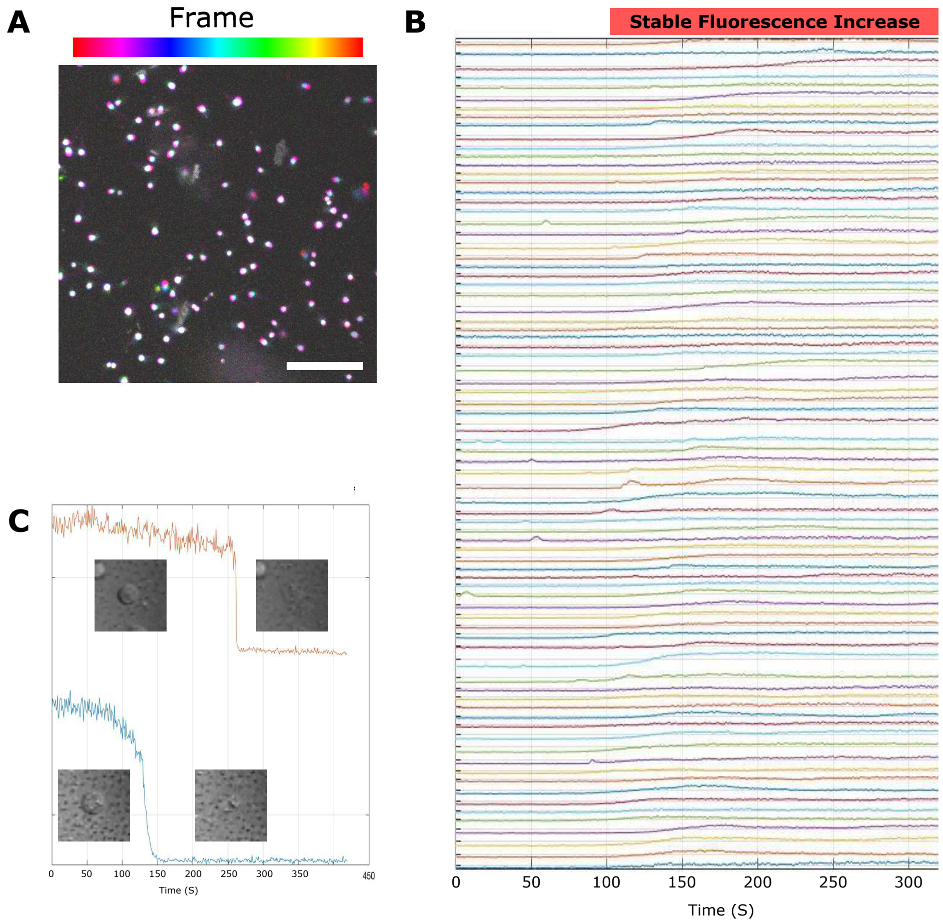


**Figure S4. Treatment with ionomycin results in prolonged stable increase in fluorescence. (A)** Temporal projection following application of 100ug/mL ionomycin. White coloration of each cell shows stable fluorescence over time series (Supplemental video 3). **(B)** Individual traces for cells from **(A)** shows maintained increase following application of the ionophore. **(C)** Traces of individual cells following large increase in laser level. Cells showed some decrease in fluorescence signal (Supplemental video 3), but loss didn't occur until the cells lysed from the exposure to high energy amounts, suggesting the indicator is highly photostable. Scale bar 50µm.

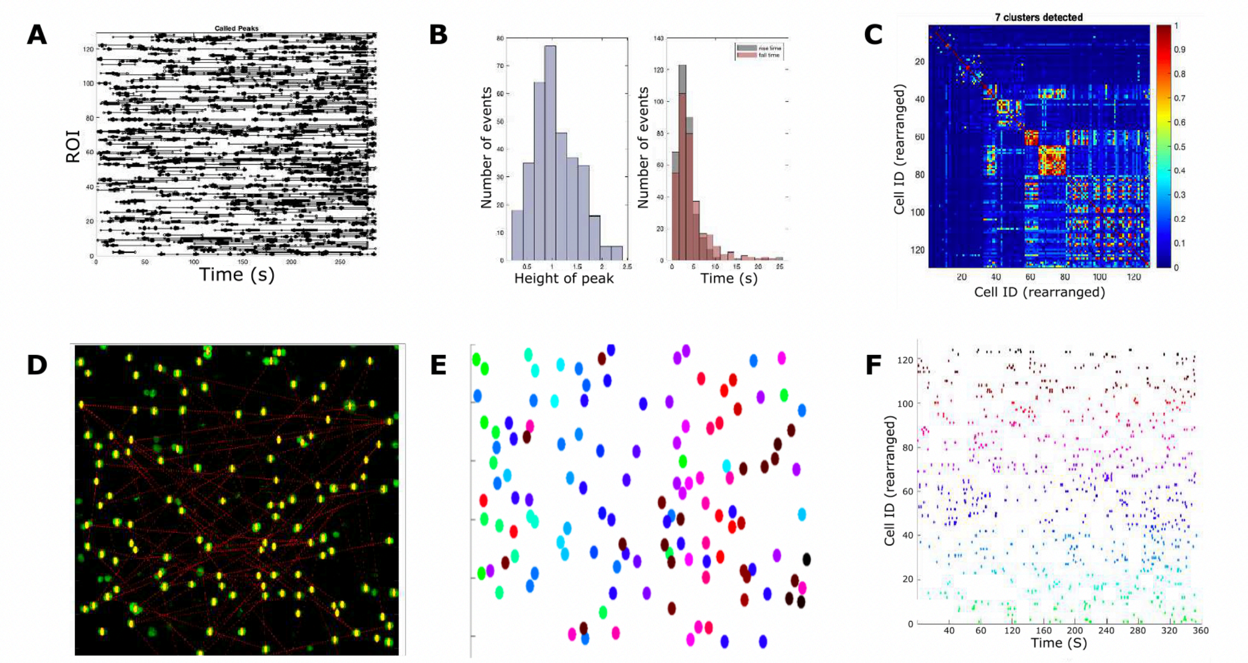


**Figure S5**. **Network activity shows limited connectivity across the field of view. (A)** Raster plot of individual events in an active spontaneously acting culture. **(B)** Histogram of basic shape characteristics of individual events shown in **(A)**. Height of peaks on left and rise (grey) and fall (red) time shown on the right. **(C)** Correlation matrix for ROIs from **(A)** finds 7 weak clusters of activity. **(D)** Node analysis does not identify any strong hubs of activity in the field of view. **(E)** ROIs in field of view color coded based on clusters from **(C)**. **(D)** Raster plot (only showing peaks) rearranged based on clusters. Clusters of activity (yellow circle) appear to occur in cells in similar locations.

**A**

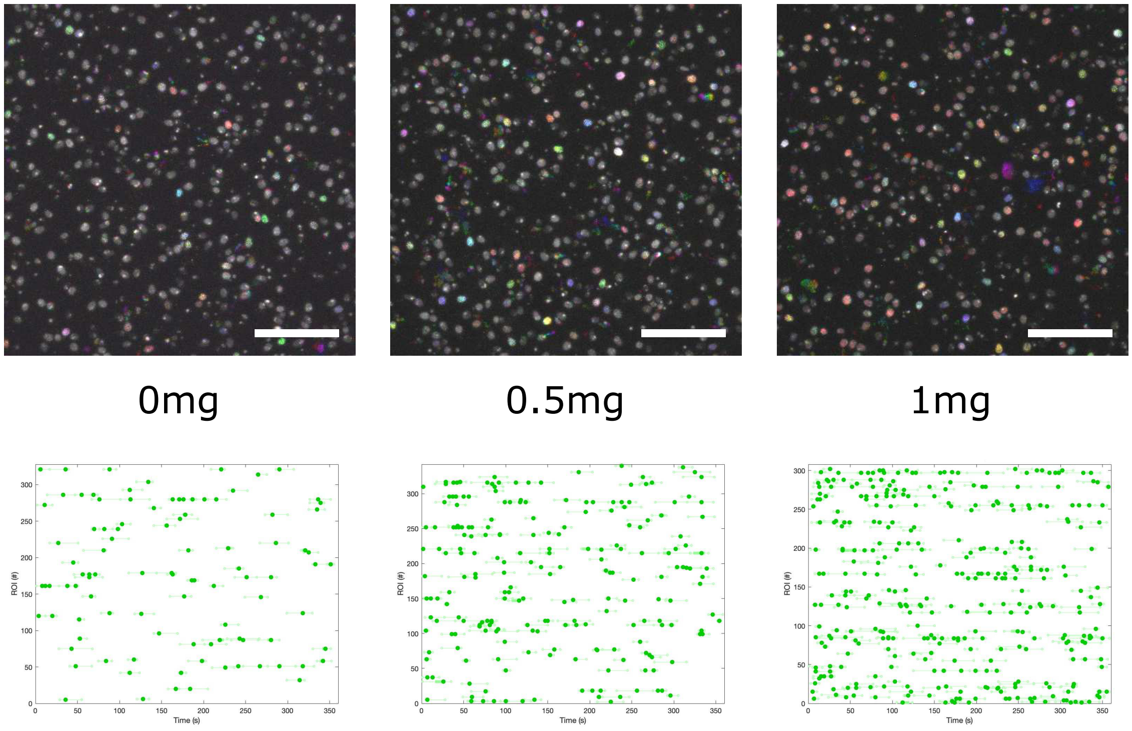


**B**

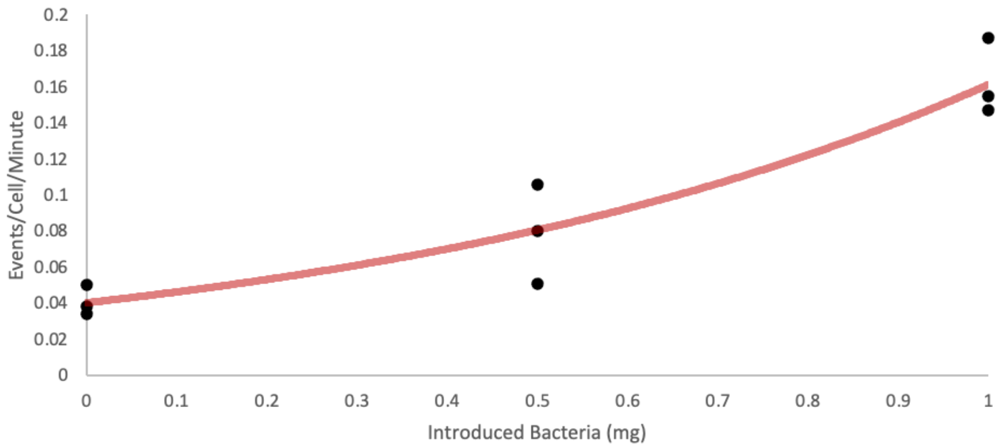


**Figure S6. Increasing bacteria added after washing increases spontaneous activity. (A)** Temporal projections (top) and raster plots (bottom) of cultures imaged after being washed with ASW and then incubated with either ASW, ASW+0.5mg *E. pacifica*, or ASW+1.0mg *E. pacifica* (Supplemental video 4). **(B)** Probability of peaks being detected in spontaneously acting cultures increases as the concentration of feeder bacteria increases. Scale bars 50µm.

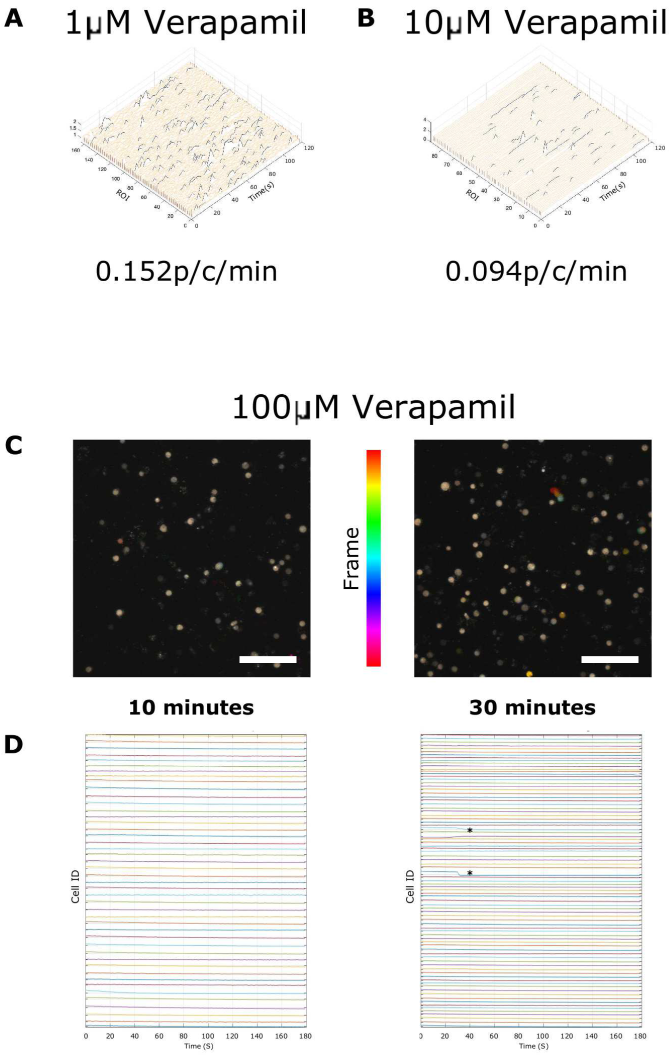


**Figure S7. Dose dependent response of cells to verapamil treatement (A)** Traces for cells incubated with 1μM verapamil and imaged for spontaneous activity (Supplemental video 5). Events occurred at a rate of 0.152 peaks/cell/min (p/c/m). **(B)** Traces for cells incubated with 10μM verapamil and imaged for spontaneous activity (Supplemental video 5). Events occurred at a rate of 0.094 peaks/cell/min (p/c/m). **(C)** Temporal projection of image sequence taken following incubation in 100uM verapamil for 10 minutes (left) and 30 minutes (right) show little activity (Supplemental video 5). Rounded morphology of cells and high base line fluorescence suggests the individual cells are not healthy. **(D)** Individual traces for cells in **(C)**. Asterisks indicate rapid decreases in cells, similar to those observed following laser ablation. Scale bars 50μm.

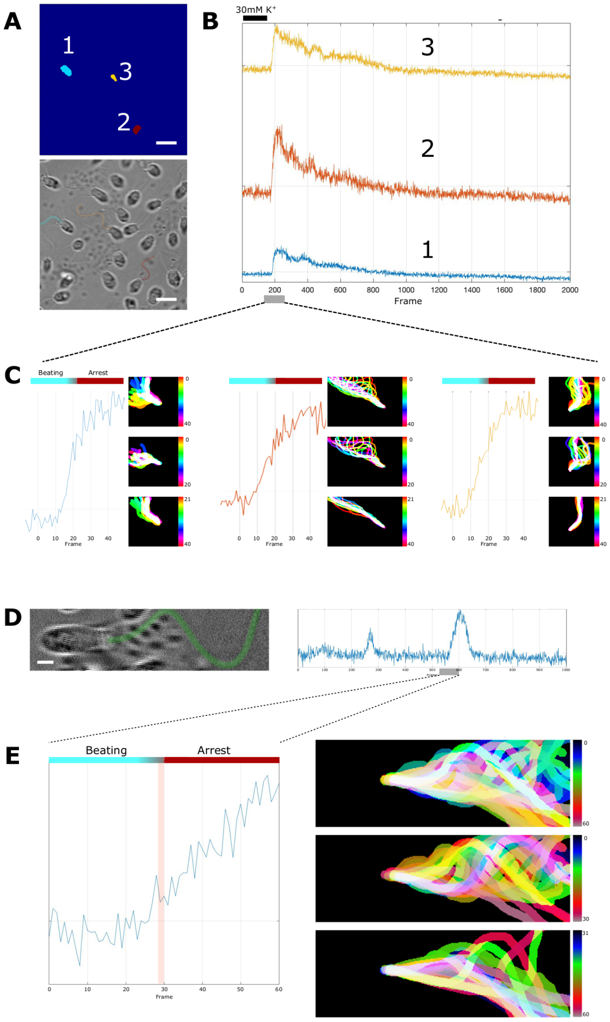


**Figure S8. Increased signal in the cell body occurs prior to ciliary arrest. (A)** ROIs chosen from video shown below based on visibility and orientation of beating plane for the flagella. **(B)** Individual traces for the three cells shown in **(A)** over the full-time course of the video. **(C)** Zoom in on the region of the traces following stimulation. The fluorescence signal begins to increase between frame 9-13 (270-390ms) of the zoomed region. Ciliary traces during the same time frame are shown next to the graph. Top shows full time span, middle shows the first 20 frames (600 ms**)**, and bottom shows last 20 frames (630-1200ms). Straightening and arrest are not observed until approximately frame 20, about 30-60ms following increased signal in the cell body. **(D)** An example cell from an unstimulated culture, where the flagella (highlighted in green) is clearly visible (Supplemental video 6). **(E)** Zoom in on the 60 frames during which a $Ca^{2+}$ transient event was occurring. Increase begins during the time from frame ~15-30. Corresponding ciliary traces are shown next to the graph showing the full sequence (top), first 30 frames (middle), and last 30 frames (bottom). Arrest does not occur until after frame 30. Scale bars 5µm (**A**) and 2µm (**B**).

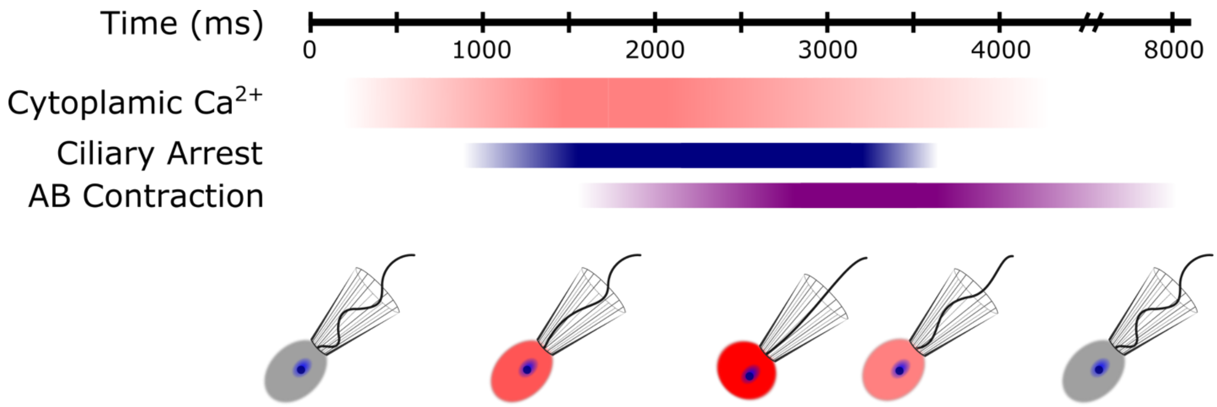


**Figue S9. General sequence of events for behavior.** Initial rise in cytoplasmic RGECO signal peaks in ~1000ms and plateaus for ~400ms before returning to baseline levels over ~2500ms. Ciliary arrest begins 600-900ms and takes ~600ms to fully arrest. Beating returns with the drop in cytoplasmic calcium levels. Apical-basal (AB) contraction begins about 500ms after the onset of ciliary arrest and continues for around 2000-3000ms before gradually returning to the initial shape.

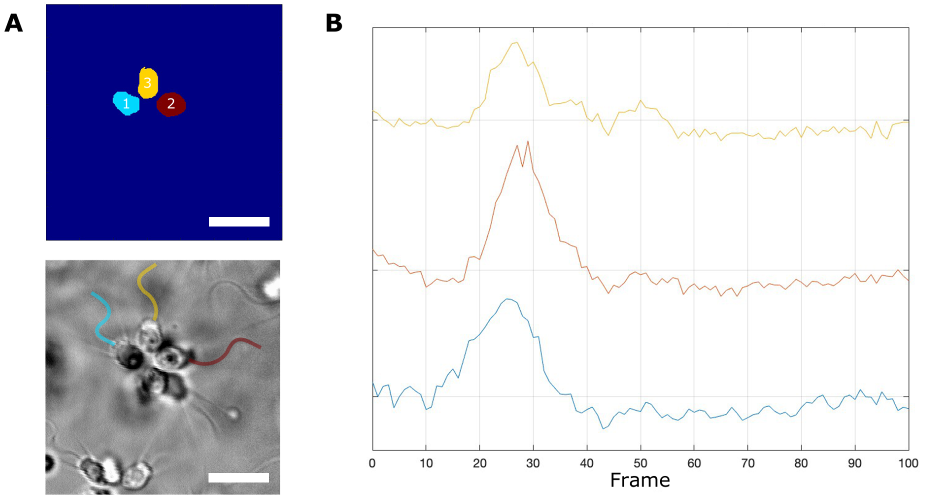


**Figure S10. Coordinated ciliary arrest in colonies occurs following initial signal in the cell body.** **(A)** Segmentation of ROIs (top) and brightfield image (bottom) of a colony undergoing a $Ca^{2+}$ transient event, with cilia highlighted. **(B)** deltaF/F0 traces for each ROI over course of video. Scale bar 10µm.

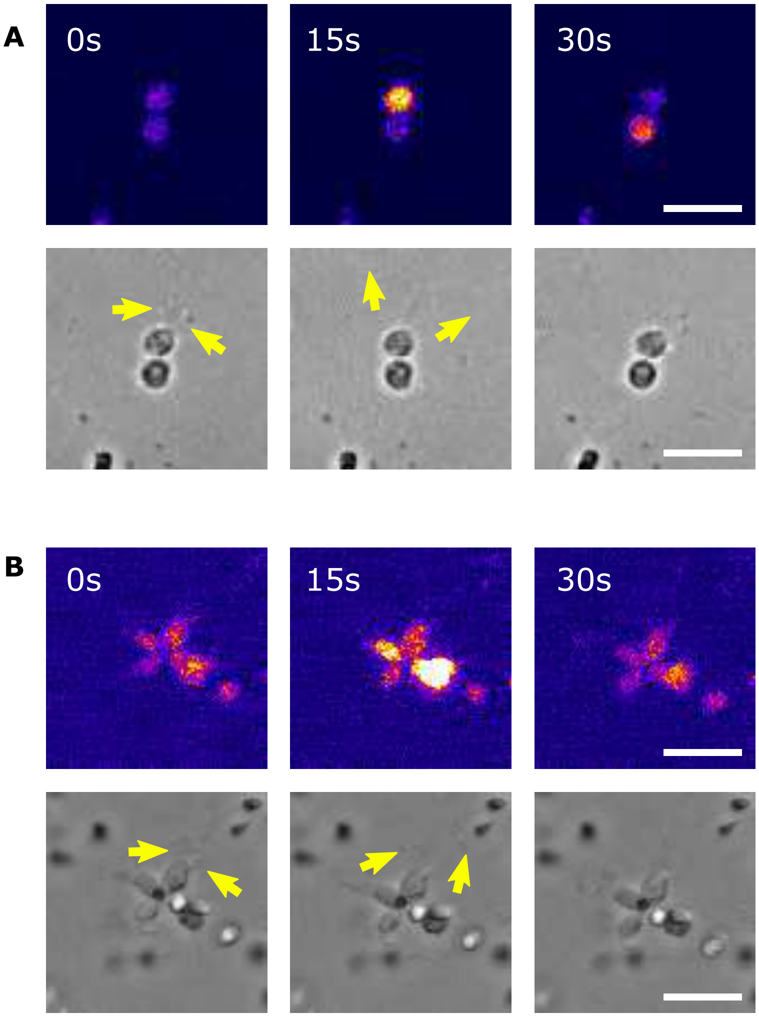


**Figure S11. Displacement of bacteria from collar in colonies. (A)** An asynchronous event in a two cell chain shows the top cell displacing the bacteria (yellow arrows) from its collar during a $Ca^{2+}$ transient (Supplemental video 7). **(B)** Bacteria (yellow arrows) visible on the collar of the cell at the top of the collar as seen to be displaced during a synchronized $Ca^{2+}$ transient (Supplemental video 7). Scale bars 10µm.

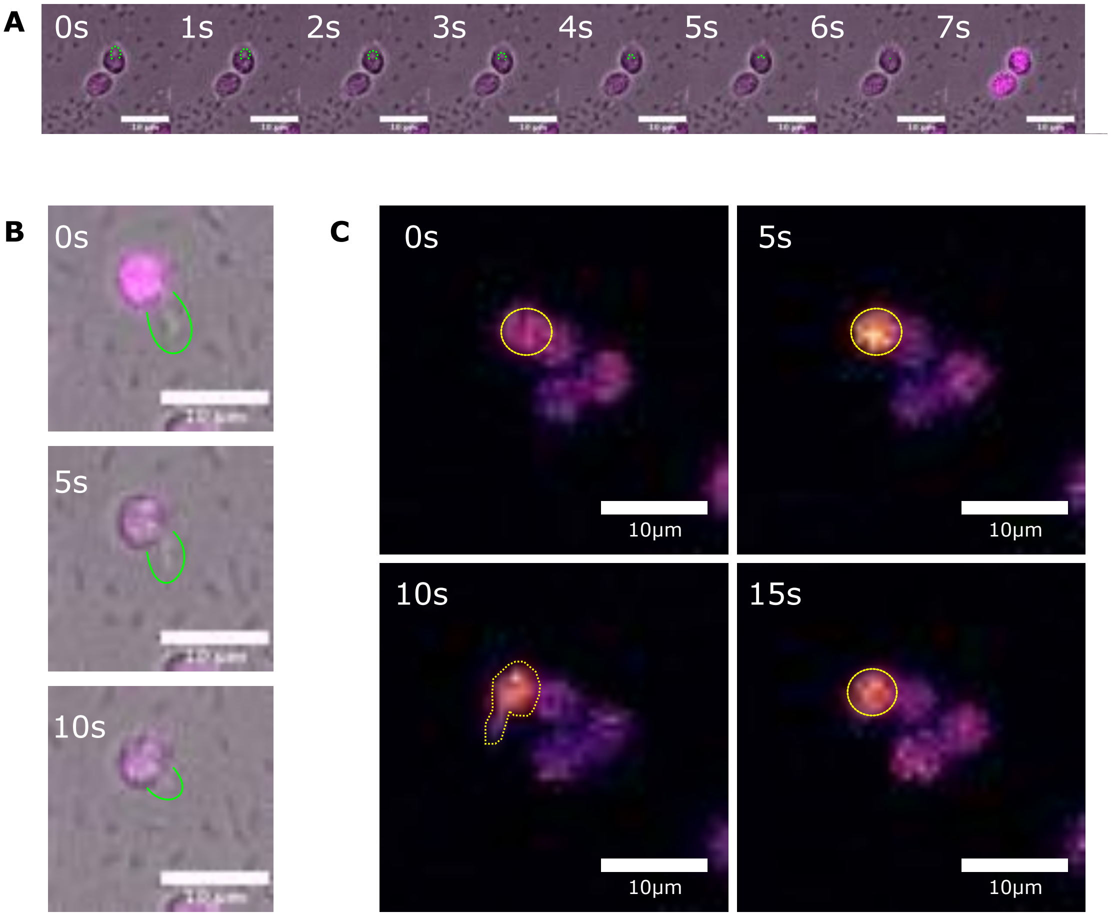


**Figure S12**. **Transients observed associated with phagocytosis. (A)** An example of a strong $Ca^{2+}$ occurring at the time the phagocytic cup (highlighted in green) is fully retracted (Supplemental video 8). **(B)** An example of an event occurring at the time that the phagocytic cup (highlighted in green) begins to retract (Supplemental video 8). **(C)** Example of an event occurring in a cell (yellow) just prior to formation of the phagocytic cup forming (bottom left panel). Increased $Ca^{2+}$ signal can be seen inside the newly formed cup and persists until retraction (Supplemental video 8).

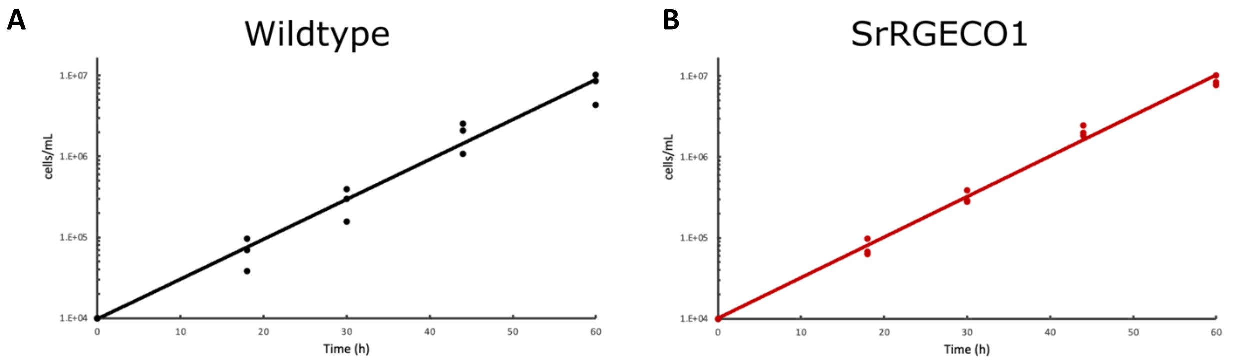


**Figure S13. Growth curves for wildtype SrEpac and SrRGECO1 are consistent. (A)** Growth curve for wildtype cultures after seeding at $10^4$ cells/mL and incubating under normal conditions. **(B)** Growth curve for SrRGECO1 cultures after seeding at $10^4$ cells/mL and incubating under normal conditions.

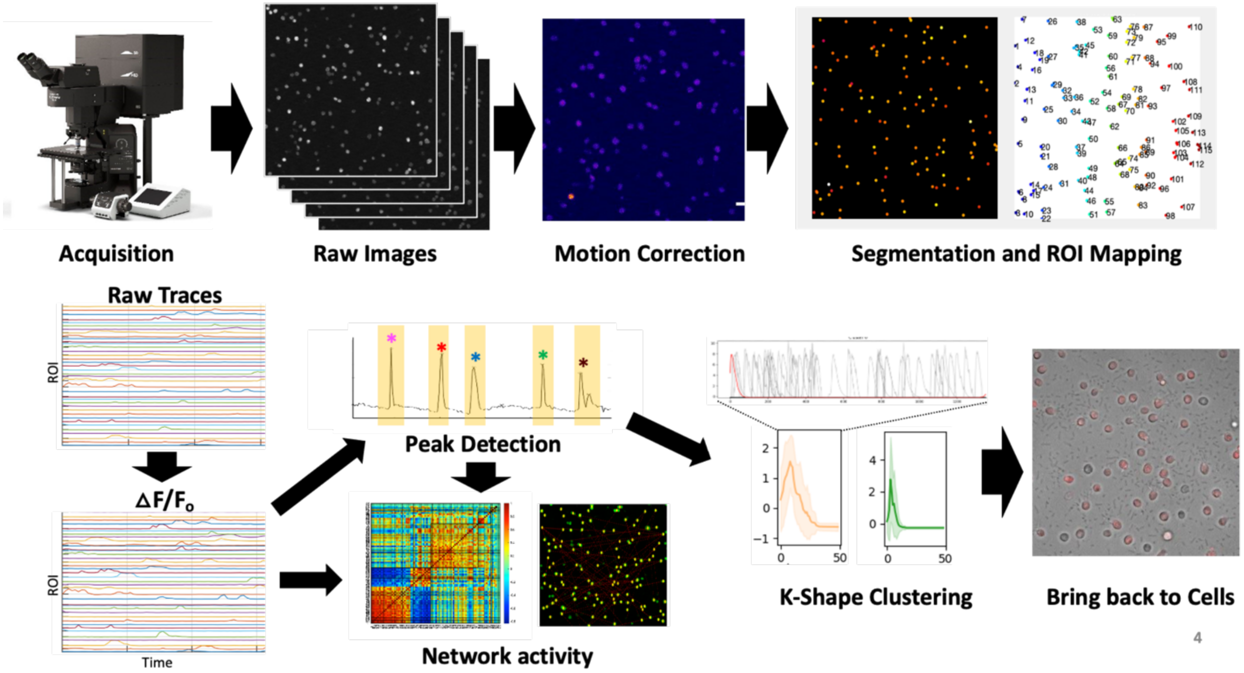


**Figure S14. Basic workflow and analysis pipeline.** Images were acquired on an Olympus FV3000RS confocal microscope using a resonance scanner with an open aperture to optimize speed. Summed projections were taken at each timepoint and processed into image sequences. Motion correction was performed using the CaImAn algorithm [8]. Segmentation was performed using either CNMF(E) [10] or active contour seeding [5]. Raw traces were extracted for each ROI and deltaF/F0 was calculated. Individual peaks were then identified. Network activity was calculated based on full traces for ROIs and individual peaks [5].

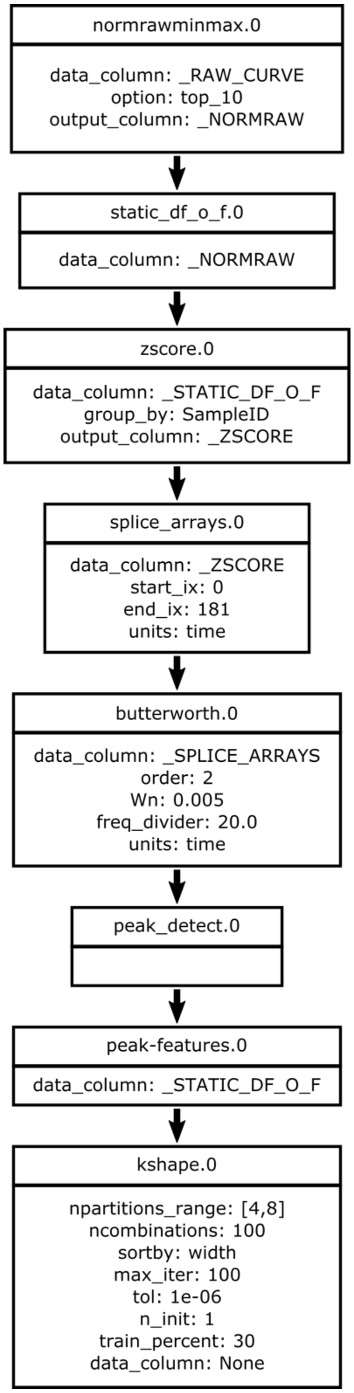


**Figure S15. k-Shape clustering analysis graph.** A graphical representation of the analysis steps and parameters used to generate the clusters in **Figure 2.** After normalizing and smoothing $dF/F_0$ curves, individual peaks were extracted. kShape clustering was performed using the parameters shown in the box, based on empirical testing of the data sets.
